# Hypoxia induced oxidative stress and endoplasmic reticulum stress promoted myocardial cell fibrosis

**DOI:** 10.1101/2023.06.24.546381

**Authors:** Zhan Jiang, Zhang Chun, Xu Guang

## Abstract

Myocardial cells, fibroblasts and vascular cells in the heart are connected by a complex matrix mainly composed of fibrillar collagen, which helps to protect the integrity and compliance of the heart structure. Previous studies have shown that hypoxia can induce myocardial hypoxia, but the mechanism is still unclear. In this study, we found that hypoxia promotes TGF beta induced collagen deposition and myocardial fibrosis by inducing Endoplasmic reticulum stress and oxidative stress in cardiomyocytes. Moreover, we also found that antioxidant drugs can effectively alleviate hypoxia induced myocardial fibrosis. Therefore, our study provides an experimental basis for the treatment of myocardial fibrosis.

## INTRODUCTION

Organ injury caused by organ ischemia-reperfusion is currently a common clinical disease, especially myocardial ischemia-reperfusion injury^[1]^. Research has shown that a major factor in myocardial injury caused by myocardial ischemia-reperfusion is hypoxia and reoxygenation^[2]^. As is well known, oxygen deficiency can induce an upregulation of hypoxia inducible factor(HIF) expression levels. HIF is the main regulator of cell response during hypoxia stress and is generally considered as the rapid response force of cell response driven by hypoxia. HIF is a heterodimeric transcription factor composed of an oxygen dependent α Subunit (1 α, 2 α or 3 α) And a constitutive expression β The subunit composition, its expression and function mainly depend on α Subunits^[3]^. Currently, the most studied area is HIF-1 α and HIF-2 α, Among them, HIF-1 α Expressed in most organs, while HIF-2 α It is expressed in specific organs. As an important hypoxia inducer, activated HIF can promote over a hundred target genes, such as vascular endothelial growth factor and glucose transporter protein^[4]^. Moreover, studies have shown that HIF is involved in the conversion of cells from oxidative phosphorylation to glycolysis. In addition, HIF can also participate in regulating fibrosis, cell death, and inflammatory responses^[5]^.

The endoplasmic reticulum is a membrane organelle with multiple functions in eukaryotic cell. Its functions mainly regulate protein folding, free calcium ion storage and nuclear lipid synthesis^[6]^. The protein folding function of endoplasmic reticulum is particularly sensitive to organ ischemia-reperfusion. Under pathological conditions, when many unfolded proteins are misfolded, stress signals will be transmitted from the endoplasmic reticulum membrane to the nucleus, thus causing a series of specific target gene transcription and protein translation levels to be down regulated. Studies have shown that HIF and endoplasmic reticulum stress (ERS) are mutually regulated^[7]^. If ischemia leads to hypoxia, which activates HIF and protects the kidneys from acute ischemic damage by regulating ERS.

Many studies have reported that hypoxia can induce endoplasmic reticulum stress response by activating HIF, thus leading to cerebral ischemia-reperfusion injury^[8]^. However, at present, it is still unknown whether the endoplasmic reticulum stress response will be activated due to hypoxia in the process of myocardial ischemia reperfusion, which will lead to myocardial fibrosis. In this study, we placed myocardial cells in a hypoxic environment to simulate the hypoxic environment of the myocardium. We found that long-term hypoxia treatment can induce myocardial cell fibrosis. Furthermore, the myocardial cells treated with hypoxia were detected. The results showed that hypoxia induced the endoplasmic reticulum stress of myocardial cells. Moreover, inhibition of endoplasmic reticulum stress can alleviate hypoxia induced myocardial fibrosis.

## Materials and Methods

### Materials

mouse monoclonal antibodies against collagen I (Sigma; C2456) or α-smooth muscle actin (αSMA, sigma; A2547) and Rabbit monoclonal antibodies against glucose regulated protein 78 (GRP78, ABclonal; A4908) or CHOP (ABclonal; A20987) were used for western blot analysis. Reactive Oxygen Species Assay Kit (Beyotime; S0033S) were used for cellular ROS measurement, NADP+/NADPH Assay Kit with WST-8 (Beyotime; S0179) were used for cellular NADPH/NADP^+^ detected. Total Glutathione Assay Kit (Beyotime; S0052) were used for cellular GSH/GSSG detected, recombinant TGF-β1 (sigma; GF346) and N-Acetylcysteine (sigma; A9165) were used as description.

### Cell culture

The cardiac muscle HL-1 cell line (ATCC, USA) were maintained in Dulbecco’s modified Eagle’s medium supplemented with 10% fetal bovine serum and antibiotics, HL-1 cell cultured under 95% air and 5% CO2 at 37°C. Culture medium was changed every 2 days.

### TGF-β1 and NAC treatment

For TGF-β1 treatment, 10 ng/mL TGF-β1 (Sigma, USA) was added into the culture medium and cells were incubated for various times; For NAC treatment, 5mM NAC (Sigma, USA) was added into the culture medium and cells were incubated for various times.

### Hypoxia treatment

For hypoxia treatment, cells were incubated in a hypoxia workstation (Baker Ruskin, UK) equilibrated with 1% O_2_/5% CO_2_/94% N_2_.

### ROS measurement

Total ROS in HL-1 cells were measured using the Reactive Oxygen Species Assay Kit (Beyotime, China), following the manufacturer’s instructions. Briefly, cells were treated as indicated and incubated with fresh medium added containing 10 mM DCFH-DA at 37°C for 30 min. The cells were then washed with PBS 3 times, cells were digested by pancreatic enzymes and collected, and then the reactive oxygen species levels were analyzed by flow cytometry (BD, USA). 1 *GSH/GSSG ration measurement*

The measurement of GSH/GSSG ratio was assessed using GSH and GSSG Assay Kit (Beyotime, China) according the manufacturer’s instructions. Briefly, 1×10^6^ cells were lysed at 4°C. GSSG in cell lysate was reduced to GSH by glutathione reductase and the GSH formed could cause a continuous reduction of 2-nitrobenzoic acid (DTNB) to 5-thio-2-nitrobenzoic acid (TNB). The concentration of TNB reflected the amount of GSH and could be measured at 420 nm spectrophotometrically. The results were normalized by protein concentration of each sample.

### NADPH/NADP^+^ ration measurement

The measurement of NADPH/NADP^+^ ratio was determined using a NADP+/NADPH Assay Kit with WST-8 (Beyotime, China) according the manufacturer’s instructions. In brief, 1×10^6^ cells were lysed by 3 frozen-thaw cycles. Lysate sample was then separated into two portions. One portion was heated at 60 °C to deplete NADP^+^ (only NADPH left) while the other portion was left on ice as unheated sample (containing both NADP^+^ and NADPH). NADP^+^ could be reduced into NADPH in the Working buffer and the NADPH formed further reduced WST-8 to formazan. The orange product (formazan) was then measured at 450 nm spectrophotometrically. The NADPH/NADP^+^ ratio was calculated using following formula: (intensity of heated sample)/ (intensity of unheated sample – intensity of heated sample). The results were normalized by protein concentration of each sample.

### RT-qPCR

Total RNA was extracted from specimens and microscopies using TRIzol (Invitrogen, USA). 1μg total RNA was synthesized using the first strand cDNA to detect the content of the target gene. QPCRs was detected with SYBR Green PCR Master Mix (Takara, Japan) reagent. Quantitative real-time PCR analysis was performed under the following conditions: 5 min at 95 °C followed by 40 cycles at 95 °C for 30 s, 55 °C for 40 s, and 72 °C for 1 min using an ABI Prism 7700 sequence detection system. Data were normalized to expression of β-actin for each experiment, The relative value of gene expression was calculated by 2-^△△CT^ method. All QPCR were three replicates, all primers were ordered from Sangon Biotech.

### Western blot analysis

Cells were collected and lysed with RIPA (Beyotime, China) containing a protease inhibitor cocktail. 20μg protein was isolated on 8-10% SDS-PAGE and transferred to nitrocellulose membrane (BioRAD). Nonspecific binding was blocked by incubation of membranes with 10% (W/V) skim milk at room temperature for 1 h. Then, the membranes were incubated with antibodies against α-SMA, collagen I (Sigma, USA); GRP78, CHOP, β-Actin (ABclonal, China) overnight at 4°C, then washed 3 times with TBST (TBS containing 1% tween 20 by volume) and incubated with secondary antibody for 1h at room temperature. Immunoreactive bands were visualized with chemiluminescence luminol reagent (Thermo Fisher). β-Actin was used to confirm equal loading of samples. Individual blots were scanned and the desired proteins quantified using densitometry.

### Statistical analysis

All data analysis used in this study were obtained from three bioinformatics replicates. GraphPad Prism9 was used for statistical analysis. The two were statistically compared using Pearson’s correlation test and Student’s *t*-test (two tailed). *P* < 0.05 was considered statistically significant.

## Results

### Hypoxia promotes TGF-β-induced collagen expression in HL-1 cells

As shown in Figure 1, cardiomyocyte HL-1 was treated with 10 ng/mL TGF-β1 for the corresponding time, the mRNA expression levels of fibrosis-related collagen I (Figure 1a) and the protein expression levels of collagen I and α-SMA (Figure 1b) were significantly upregulated in HL-1 cardiac myocytes. This indicates the successful construction of an in vitro model for TGF-β-induced fibrosis in HL-1 cells. Surprisingly, we found that hypoxia treatment further increased the mRNA expression levels of collagen I induced by TGF-β1 (Figure 1c), as well as the protein expression levels of collagen I and α-SMA (Figure 1d). These results suggest that hypoxia promotes fibrosis progression.

**Fig 1.**
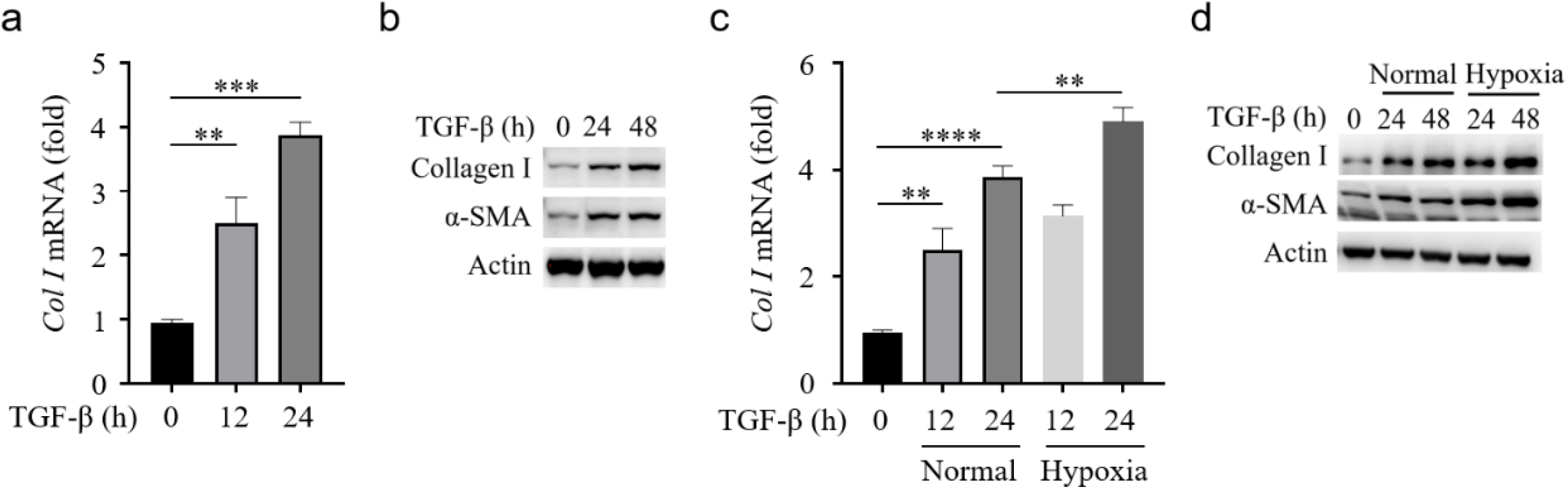
Expression of TGF-β1 and α-SMA in HL-1 cells after hypoxia. The mRNA and protein levels of collagen I were detected in HL-1 cells at the corresponding time of 10μg/ml TGF-β treatment (a, b), Effect of hypoxia on expression of collagenI and α-SMA in HL-1 cells (c, d). Data shown are represented as the means ± S.E.M. from three separate experiments. **P<0.01 vs. control, ****P<0.0001 vs. control.

### Hypoxia induces endoplasmic reticulum stress in HL-1 cells

Previous studies have shown that endoplasmic reticulum (ER) stress is involved in myocardial fibrosis^[9]^. We investigated whether hypoxia induces ER stress in cardiac myocytes. As shown in Figure 2, under hypoxic conditions, the mRNA expression levels of the ER chaperone protein GRP78 (Figure 2a) and the ER stress-related protein CHOP (Figure 2b) were significantly upregulated in cardiac myocytes HL-1 at the corresponding time points. Additionally, the protein expression levels of GRP78 (Figure 2a) and CHOP (Figure 2b) were significantly increased, indicating that hypoxia induces ER stress in cardiac myocytes HL-1.

**Fig 2.**
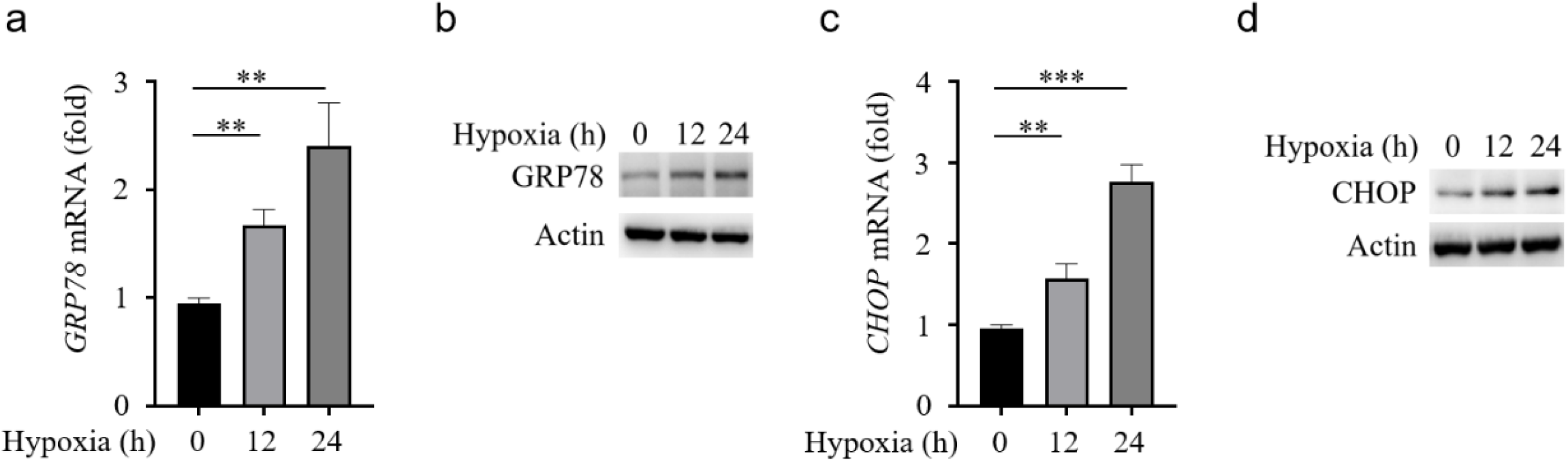
Hypoxia induce ER stress in HL-1 cells. Expression of GRP78 after different intermittent hypoxia times in HL-1 cells (a, b). Expression of CHOP after different intermittent hypoxia times in HL-1 cells (c, d). Data shown are represented as the means ± S.E.M. from three se parate experiments. **P<0.01 vs. control, ***P<0.001 vs. control.

### Hypoxia induces oxidative stress in HL-1 cells

Research has shown that reactive oxygen species (ROS) is an important factor leading to endoplasmic reticulum (ER) stress^[10]^. In order to investigate the potential mechanism of hypoxia-induced ER stress, we examined the effects of hypoxia on ROS levels in cells. As shown in Figure 3, the levels of intracellular ROS in HL-1 cardiac cells significantly increased under hypoxia treatment for the corresponding duration (Figure 3a, 3b). At the same time, the ratio of the antioxidant glutathione (GSH) to the oxidized form of glutathione (GSSG) significantly decreased (Figure 3c), and the level of intracellular reducing agent NADPH also significantly decreased (Figure 3d). These findings suggest that hypoxia may induce ER stress in HL-1 cardiac cells through ROS generation.

**Fig 3.**
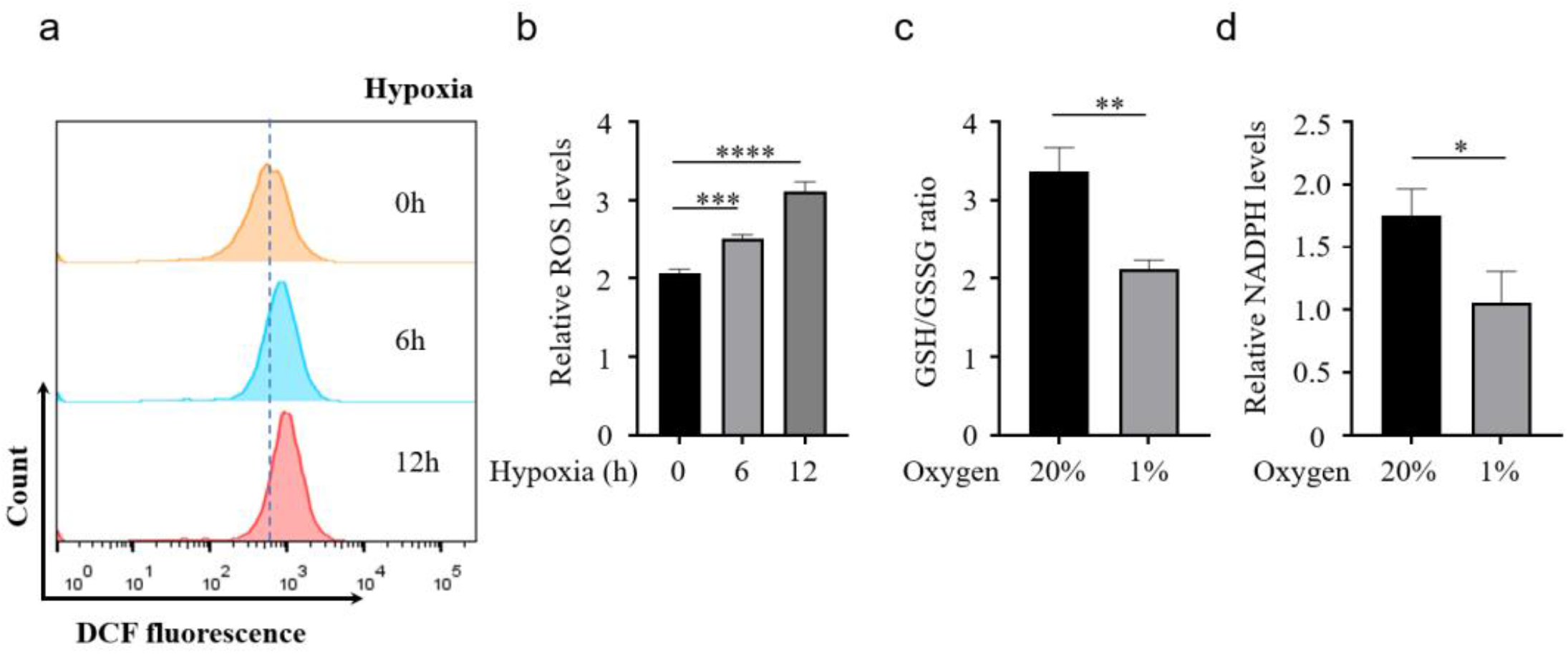
Hypoxia increase cellular levels of ROS and decrease the levels of GSH and NADPH. The ROS levels after different intermittent hypoxia times in HL-1 cells (a, b). Effect of hypoxia on GSH/GSSG ratio in HL-1 cells (c). Effect of hypoxia on NADPH levels in HL-1 cells. Data shown are represented as the means ± S.E.M. from three separate experiments. *P<0.05 vs. control, **P<0.01 vs. control, ***P<0.001 vs. control.

### Antioxidant drugs NAC reduce Hypoxia-induced endoplasmic reticulum stress in HL-1 cells

Our results show that hypoxia may induce endoplasmic reticulum stress in cardiomyocytes through ROS, further promoting the process of myocardial fibrosis. To investigate the role of antioxidants in myocardial fibrosis, we examined the effect of the antioxidant N-Acetylcysteine (NAC) on hypoxia-induced endoplasmic reticulum stress in cardiomyocytes. As shown in Figure 4, 5mM NAC, an antioxidant, significantly reduces hypoxia-induced ROS production in cardiomyocytes (Figure 4a, 4b). Additionally, we found that 5mM NAC significantly reduces the protein expression levels of the endoplasmic reticulum chaperone protein GRP78 and the endoplasmic reticulum stress-related protein CHOP induced by 10 ng/mL TGF-β1 treatment for 24 hours in cardiomyocytes (Figure 4c). These results indicate that NAC, an antioxidant, significantly inhibits hypoxia-induced endoplasmic reticulum stress.

**Fig 4.**
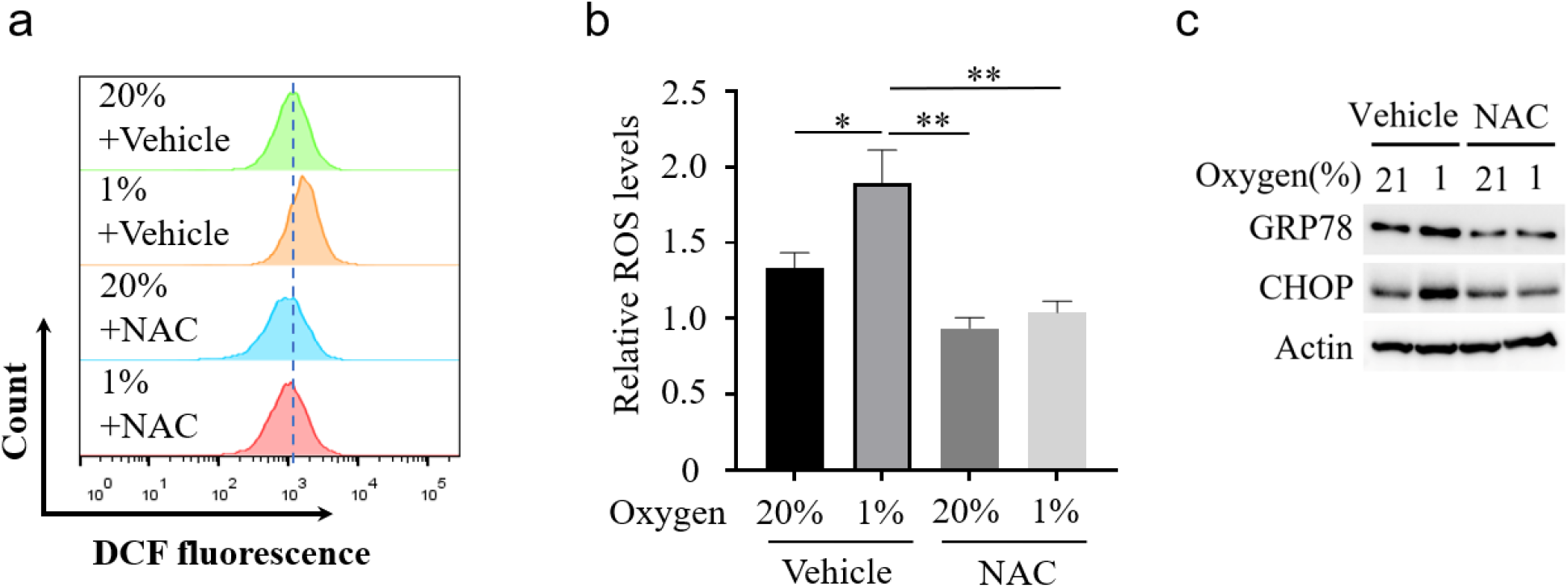
Antioxidant drugs NAC reduce cellular levels of ROS, and reduce GRP78 and CHOP expression in HL-1 cells. ROS levels in hypoxia-induced HL-1 cells were measured after 5mM NAC treatment (a, b). Effect of 5mM NAC on GRP78 and CHOP expression in HL-1 cells (c). Data shown are represented as the means ± S.E.M. from three separate experiments. *P<0.05 vs. control, **P<0.01 vs. control.

**Fig 5.**
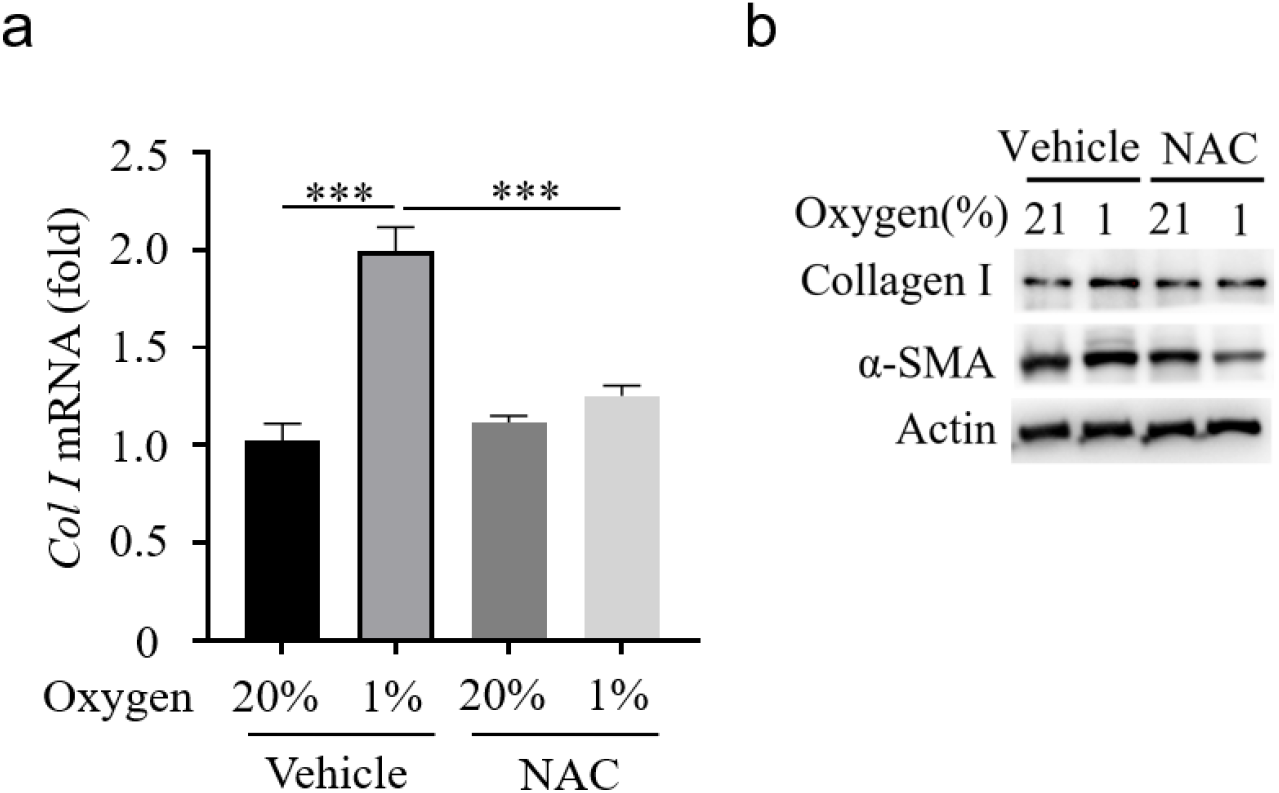
Antioxidant drugs NAC reduce α-SMA and collagen expression in HL-1 cells. The mRNA levels in hypoxia-induced HL-1 cells were measured after 5mM NAC treatment (a). The protein levels of collagen I and α-SMA in hypoxia-induced HL-1 cells were measured after 5mM NAC treatment (b). Data shown are represented as the means ± S.E.M. from three separate experiments. ***P<0.001 vs. control.

### Antioxidant drugs NAC decrease collagen expression in HL-1 cells

To further explore the role of antioxidants in myocardial fibrosis, we examined the effects of the antioxidant NAC on collagen production in hypoxia-induced myocardial cells. As shown in Figure 5, when HL-1 myocardial cells were treated with 10 ng/mL of TGF-β1 for 24 hours, the antioxidant NAC at a concentration of 5 mM significantly reduced collagen I mRNA levels in the myocardial cells (Figure 5a).

Additionally, the antioxidant NAC at a concentration of 5 mM significantly decreased the protein expression levels of collagen I and α-SMA in the myocardial cells (Figure 5b). These results indicate that the antioxidant NAC can significantly inhibit the progression of myocardial fibrosis.

## DISSUSSION

Myocardial infarction is a common heart disease, and research has shown that myocardial infarction is often accompanied by the appearance of fibrosis factors, such as hypoxia and TGF-β. As is well known, TGF-β is an important factor promoting myocardial fibrosis^[11]^. During this process, TGF-β Can promote α-Expression of SMA. However, hypoxia affects TGF-β the regulation and mechanism of inducing myocardial fibrosis are still unclear. In this study, we found that hypoxia can promote TGF-β There is large myocardial cell fibrosis. In terms of mechanism, we found that hypoxia may participate in the regulation of TGF by inducing oxidative stress and Endoplasmic reticulum stress-β Induced myocardial cell fibers.

In addition, studies still show that the increase of hypoxia time can promote α-Expression of α-SMA. At the same time, some studies have shown that hypoxia can activate ERK, JNK and p38 signaling pathways and strengthen myocardial fibrosis. Studies have shown that Endoplasmic reticulum stress can strengthen myocardial injury caused by myocardial ischemia and reperfusion. In addition, studies have also shown that Endoplasmic reticulum stress can strengthen myocardial fibrosis. Therefore, these studies just suggest that hypoxia may enhance TGF by inducing Endoplasmic reticulum stress-β Induced myocardial fibrosis. At the same time, we also realize that there are still many mechanisms of hypoxia regulating myocardial fibrosis to be further explored.

## ACKNOWLEDGEMENTS

No

## AUTHOR CONTRIBUTIONS

Z.J., Z.C. conceived the project and designed the research; Z.J. carried out most of the experiments;

Z.J. analyzed the data; Z.J., X.G. wrote the manuscript.

DECLARATION OF INTERESTS

The authors declare no competing interests.

## Reference

1. Heusch G. Myocardial ischaemia-reperfusion injury and cardioprotection in perspective. Nat Rev Cardiol 2020; 17(12):773–789.

2. Ruan W, Ma X, Bang IH, Liang Y, Muehlschlegel JD, Tsai KL, et al. The Hypoxia-Adenosine Link during Myocardial Ischemia-Reperfusion Injury. Biomedicines 2022; 10(8).

3. Semenza GL. HIF-1 and mechanisms of hypoxia sensing. Curr Opin Cell Biol 2001; 13(2):167–171.

4. Kierans SJ, Taylor CT. Regulation of glycolysis by the hypoxia-inducible factor (HIF): implications for cellular physiology. J Physiol 2021; 599(1):23–37.

5. Xiong A, Liu Y. Targeting Hypoxia Inducible Factors-1alpha As a Novel Therapy in Fibrosis. Front Pharmacol 2017; 8:326.

6. Schwarz DS, Blower MD. The endoplasmic reticulum: structure, function and response to cellular signaling. Cell Mol Life Sci 2016; 73(1):79–94.

7. Chipurupalli S, Kannan E, Tergaonkar V, D’Andrea R, Robinson N. Hypoxia Induced ER Stress Response as an Adaptive Mechanism in Cancer. Int J Mol Sci 2019; 20(3).

8. Delbrel E, Soumare A, Naguez A, Label R, Bernard O, Bruhat A, et al. HIF-1alpha triggers ER stress and CHOP-mediated apoptosis in alveolar epithelial cells, a key event in pulmonary fibrosis. Sci Rep 2018; 8(1):17939.

9. Luo T, Kim JK, Chen B, Abdel-Latif A, Kitakaze M, Yan L. Attenuation of ER stress prevents post-infarction-induced cardiac rupture and remodeling by modulating both cardiac apoptosis and fibrosis. Chem Biol Interact 2015; 225:90–98.

10. Zeeshan HM, Lee GH, Kim HR, Chae HJ. Endoplasmic Reticulum Stress and Associated ROS. Int J Mol Sci 2016; 17(3):327.

11. Yan Z, Shen D, Liao J, Zhang Y, Chen Y, Shi G, et al. Hypoxia Suppresses TGF-B1-Induced Cardiac Myocyte Myofibroblast Transformation by Inhibiting Smad2/3 and Rhoa Signaling Pathways. Cell Physiol Biochem 2018; 45(1):250–257.

